# Activity of EGFR transmembrane region variants indicates specific transmembrane dimers are not required for EGFR activity

**DOI:** 10.1101/2022.09.08.507182

**Authors:** Foteini Bartzoka, Monica Gonzalez-Magaldi, Patrick O. Byrne, Nicole I. Callery, Kalina Hristova, Daniel J. Leahy

**Affiliations:** Department of Molecular Biosciences, University of Texas at Austin, 100 E. 24^th^ St. Stop A5000, Austin, TX 78712; Department of Materials Science and Engineering, Johns Hopkins University, 3400 Charles St., Baltimore, MD 21218

**Keywords:** EGFR, transmembrane domain, phosphotyrosine signaling, Erk, Akt

## Abstract

The Epidermal Growth Factor Receptor (EGFR) is a Receptor Tyrosine Kinase that mediates cell proliferation and differentiation events during development and maintenance of complex organisms. Formation of specific, ligand-dependent EGFR dimers is a key step in stimulating EGFR signaling, and crystal structures of active, dimeric forms of isolated EGFR extracellular regions and kinase domains have revealed much about how dimer interactions regulate EGFR activity. The nature and role of the transmembrane region in regulating EGFR activity remains less clear, however. Proposed roles for the transmembrane region range from nonspecific but energetically favorable interactions to specific transmembrane dimer conformations being associated with active, inactive, or activity-modulated states of EGFR. To investigate the role of specific transmembrane dimers in modulating EGFR activity we generated thirteen EGFR variants with altered transmembrane sequences designed to favor or disfavor specific types of transmembrane region interactions. We show using FRET microscopy that EGFR transmembrane regions have an intrinsic propensity to associate in mammalian cell membranes that is counteracted by the extracellular region. We show using cell-based assays that each of the EGFR transmembrane variants except the Neu variant, which results in constitutive receptor phosphorylation, is able to autophosphorylate and stimulate phosphorylation of downstream effectors Erk and Akt. Our results indicate that many transmembrane sequences, including polyleucine, are compatible with EGFR activity and provide no evidence for specific transmembrane dimers regulating EGFR function.

## Introduction

The Epidermal Growth Factor Receptor (EGFR) was the first known example of a Receptor Tyrosine Kinase (RTK), a family of receptors typified by an extracellular ligand-binding region, a single membrane-spanning region, and an intracellular tyrosine kinase whose activity is stimulated by ligand binding (1). The human genome contains 58 RTKs that assort into 20 classes based on extracellular region homology (1). In addition to EGFR, specific RTK classes include receptors for Vascular Endothelial Growth Factor (VEGFR), Fibroblast Growth Factor (FGFR), and Insulin (InsR). RTKs are thought to signal through formation of ligand-dependent dimers with increased kinase activity (2, 3), but the presence of dimers of EGFR and other RTKs in the absence of ligand (4–6) suggests that dimerization *per se* may not be the activating signal and that formation of specific dimer conformations is needed to initiate signaling (7, 8).

Four EGFR homologs are present in humans including HER2/ErbB2, overexpression of which drives ~20% of human breast cancers (9). Seven EGFR ligands have been identified, including high-affinity ligands Epidermal Growth Factor (EGF) and Transforming Growth Factor-a (TGFa) (10). Characteristic of RTKs, ligand binding to the EGFR extracellular region stimulates its intracellular kinase activity (11–15), which in turn leads to receptor autophosphorylation, recruitment of adaptor proteins, and activation of downstream signaling effectors including the Erk and Akt kinases (16). EGFR signaling stimulates cell proliferation and differentiation and plays essential roles in the development and maintenance of animal tissues (17). Abnormal EGFR signaling is associated with many human cancers (17), and in the last two decades EGFR and its homologs have become targets of several successful anticancer therapies (18, 19).

Crystal structures of isolated extracellular and kinase regions of EGFR and its homologs in active and inactive states have revealed much about how specific receptor-ligand and receptor-receptor interactions regulate EGFR activity (13–15, 20–24), but the nature and role of specific EGFR transmembrane (TM) region conformations in active and inactive states is less clear. EGFR TM regions interact in a TOXCAT assay (25) but do not dimerize strongly in micelles (26), and earlier authors proposed both weak and passive as well as strong and specific roles for the TM region in EGFR signaling (27–30). More recent authors have noted that the EGFR and HER2 TM regions contain two Glycine-x-x-x-Glycine (GxxxG) motifs, one in the N-terminal portion and another near the C-terminus. GxxxG motifs are characterized by glycine or another small amino acid separated by 3 amino acids, which aligns glycines on one side of a transmembrane alpha helix and creates a notch favorable for homotypic TM interactions (31, 32). Indeed, NMR structures of an EGFR TM region homodimer and an EGFR/HER2 TM region heterodimer show dimer interactions mediated by the N-terminal GxxxG regions (33, 34). GxxxG motifs are not conserved in all RTK TM regions, however, including those of VEGFR and InsR.

X-ray crystallographic and crosslinking studies strongly imply that EGFR TM regions interact when EGFR is activated by ligand (35), and several types of EGFR TM interactions have been proposed. Studies using the ToxR system and synthetic peptides led to conclusion that EGFR uses the C-terminal GxxxG for homodimeric interactions and the N-terminal GxxxG for heterodimeric interactions with HER2 (36). Computational, NMR, and phosphorylation studies have suggested that TM interactions mediated by the N-terminal and C-terminal GxxxG motifs reflect active and inactive states, respectively (33, 37). More recently, crosslinking studies and computational analyses have also been interpreted to indicate different EGFR TM dimer conformations depending on whether EGF or TGFa is bound and that these different TM dimers lead to different signaling outcomes, or biased agonism (38). Kinetic-dependent explanations for biased agonism have also been proposed (39).

To investigate the role of the EGFR TM region in modulating EGFR signaling we performed both biophysical and cell-based assays. Using Quantitative Imaging FRET (QI-FRET) (40), in which proteins labeled with a suitable FRET pair are co-transfected into cells, we demonstrated an intrinsic propensity of EGFR TM regions to interact in mammalian cell membranes. Addition of the EGFR extracellular region to the TM inhibited this interaction in the absence of ligand. We then created a panel of thirteen EGFR variants with altered TM sequences designed to promote or disrupt specific types of TM interactions. These variants include TM regions in which the N-terminal GxxxG region has been removed by substitution with isoleucines (33); TM regions from human HER2/ErbB2 and native and oncogenic forms of the rat HER2/ErbB2 TM; the constitutive Glycophorin A TM dimer and a dimerization-impaired Glycophorin A TM variant (41); TM regions from human VEGFR and InsR, which do not contain GxxxG motifs; a 23-amino acid polyleucine sequence; and polyleucine TMs with GxxxG motifs inserted at N-terminal, middle, and C-terminal positions. Except for the oncogenic rat ErbB2 TM (Neu/V664E), which was constitutively phosphorylated and able to activate downstream effectors, each of the EGFR TM variants responded to stimulation with EGF or TGFα with increased autophosphorylation and activation of the downstream effector kinases Erk and Akt, similar to the wild-type. These results show that the EGFR extracellular region counteracts the EGFR TM region’s intrinsic propensity to interact with itself and that specific EGFR TM dimer structures are not required for ligand-dependent regulation of EGFR activity.

## Results

We performed FRET microscopy to determine if the isolated EGFR TM region self-associates in mammalian cell membranes. Expression plasmids encoding the EGFR signal sequence followed directly by the TM region and a C-terminal label of either Enhanced Yellow Fluorescence Protein (EYFP) or mCherry, which constitute a suitable FRET donor-acceptor pair, were cotransfected in Chinese Hamster Ovary (CHO) cells (Figure 1A). QI-FRET was used to measure receptor concentration, which varied up to 10-fold in different cells, and FRET efficiency between labeled TM regions as a function of TM region concentration in membrane-derived vesicles (Figure 1B) (40, 42). For comparison, QI-FRET was also performed on the EGFR extracellular region plus TM followed by either EYFP or mCherry in both the presence and absence of EGF (Figure 1B). Kd values for each interaction are shown in Table S1. These data show that the EGFR TM has an intrinsic propensity to self-associate in mammalian cell membranes that is counteracted by the presence of the EGFR extracellular region. The antagonistic effect of the EGFR extracellular region on TM association is relieved by addition of ligand (Figure 1B), presumably through formation of ligand-dependent EGFR extracellular region dimers observed in crystal structures (13, 14).

**Figure 1.**
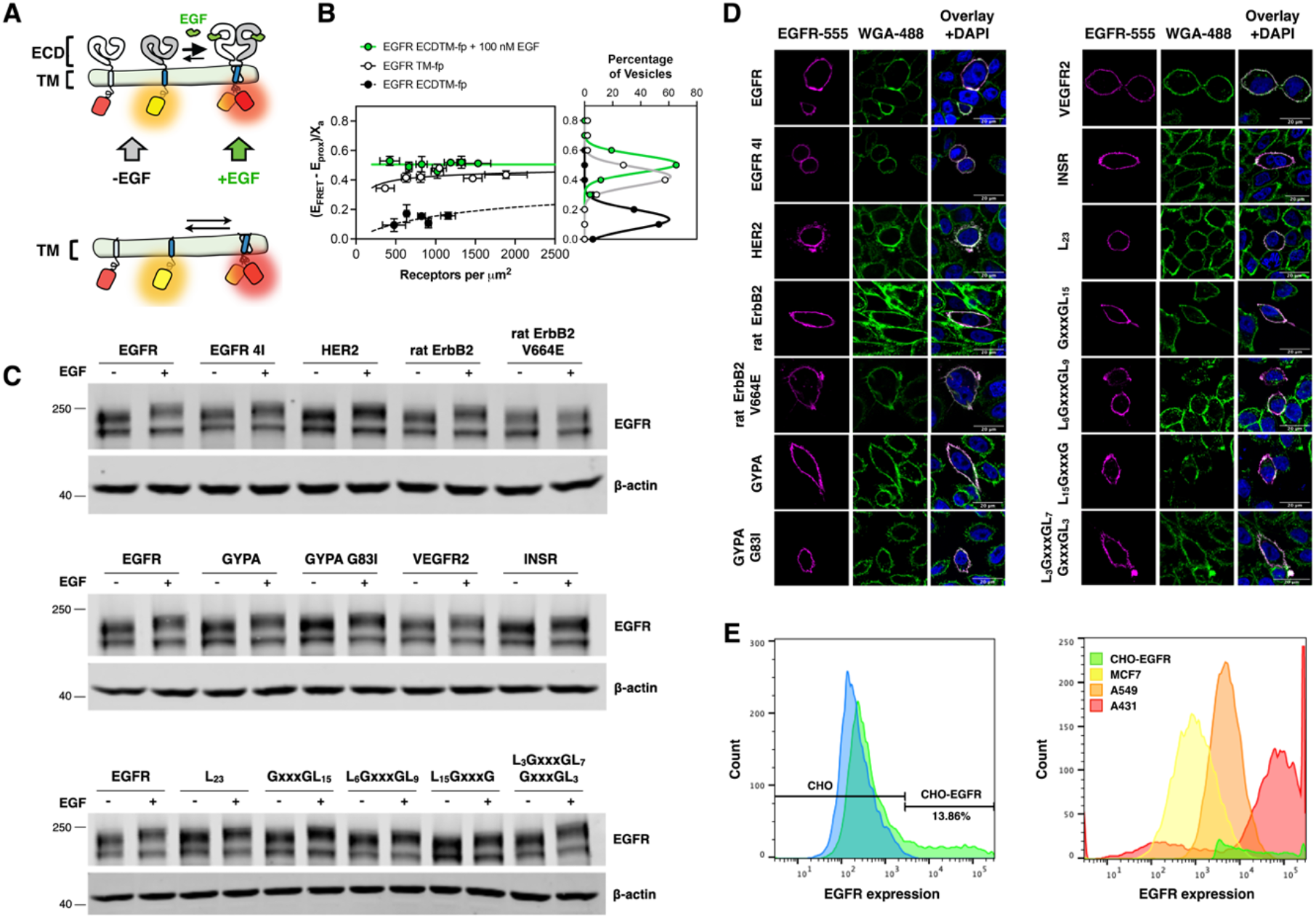
All EGFR TM variants express well and traffic to the surface in CHO cells. (A) Cartoon representation of fluorescent-protein-linked EGFR ECD-TM (top) and EGFR TM (bottom). (B) QI-FRET analysis of EGFR TM region only (EGFR TM-fp) and EGFR extracellular-TM region (EGFR ECDTM-fp) in the presence and absence of EGF. The plot on the left shows the FRET efficiency as a function of receptor concentration. The FRET efficiency of the EGFR TM region at high concentrations approaches that of the EGFR ECDTM region in the presence of ligand and is higher than the FRET efficiency of EGFR ECDTM in the absence of ligand at all concentrations. Points correspond to the average values for at least three vesicles. Error bars show the standard error in *y* and the standard deviation in *x.* The solid and dashed lines depict the lines of best fit to a monomer/dimer equilibrium model. The plot on the right shows a binned histogram of FRET values for each variant. Solid lines show the lines of best fit to a gaussian distribution. The *y*-axes for both plots are identical. (C) Anti-EGFR Western blot analysis of CHO cells transiently transfected with EGFR TM variants and treated with or without 100 nM EGF. Each panel contains wild-type EGFR for comparison and a blot for β-actin as a loading control. (D) Confocal microscopy images of transiently transfected non-permeabilized CHO cells. Receptor expression is shown in purple and cell membranes are shown in green. Scale bars represent 20 μm. Experiments were performed in triplicate and representative western blots and confocal images are shown. (E) Flow cytometry analysis of CHO-EGFR, A431, A549 and MCF7 cells expressing different levels of EGFR. Left panel shows non-transfected CHO cells in blue and CHO-EGFR transfected cells in green and transfection efficiency is shown in the histogram. Right panel shows the comparison of transiently transfected CHO-EGFR cells with cancer cell lines endogenously expressing varying levels of EGFR.

Given the intrinsic propensity of EGFR TM regions to self-associate in mammalian cell membranes, we next investigated whether specific TM dimer interactions influence EGFR signaling. DNA sequences encoding wild-type human EGFR and 13 EGFR variants with altered transmembrane (TM) region sequences were cloned into the mammalian expression vector pSSX (43). The TM variants were designed to favor or disfavor specific TM interactions as noted above, and their sequences are listed in Table 1. To aid visualization, each EGFR variant included a C-terminal Yellow Fluorescent Protein tag, which does not interfere with EGFR function (7).

**Table 1.**
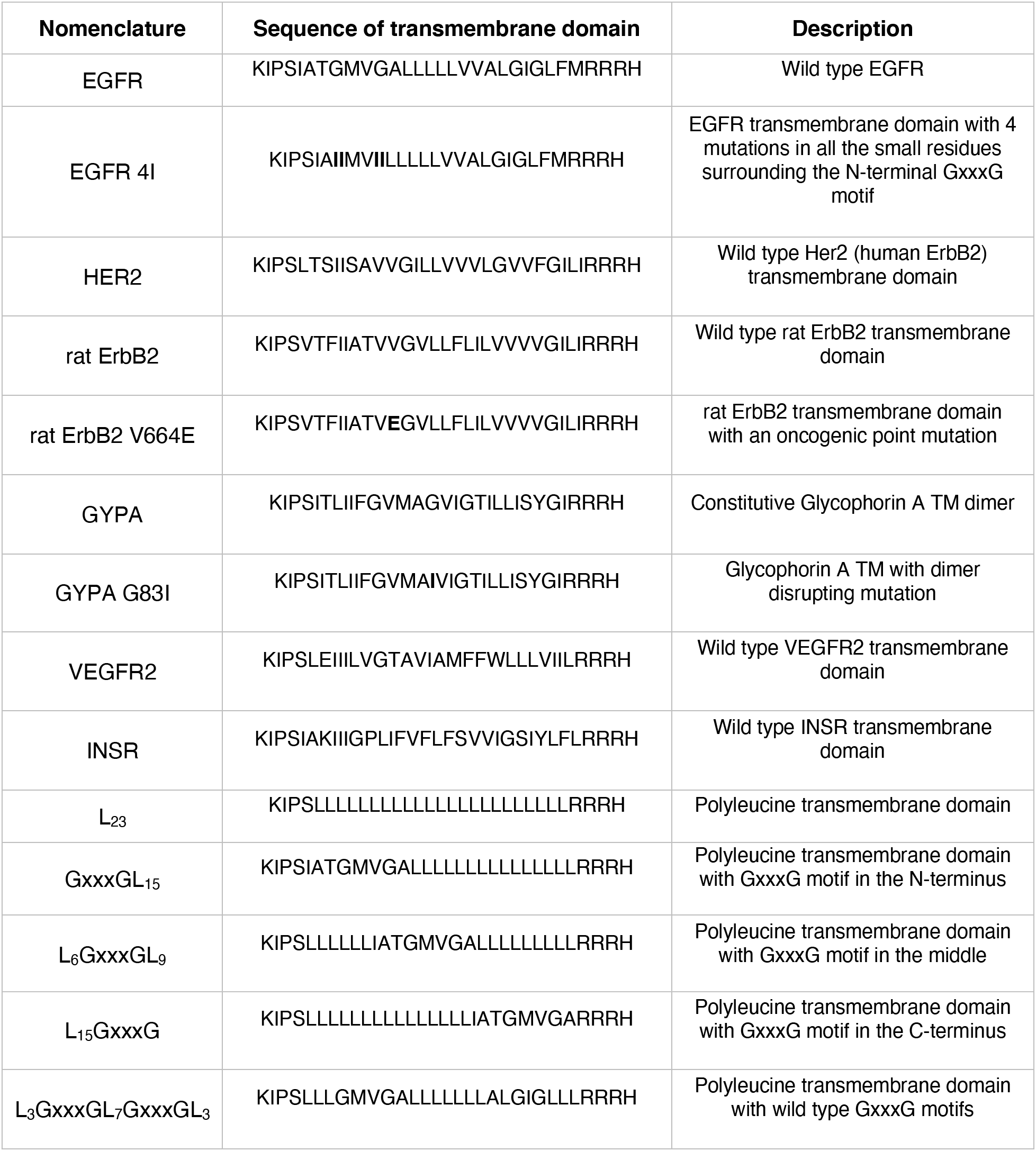
Amino-acid sequences of transmembrane domain variants.

Wild-type EGFR and each of the EGFR TM variants were transiently expressed in Chinese Hamster Ovary (CHO) cells, which do not express endogenous EGFR. All TM variants trafficked to the cell surface and expressed at levels comparable to that of native EGFR (Figure 1C-D). Transfection efficiencies were typically 12-17% as judged by immunofluorescence and fluorescence-activated cell sorting (Figure 1E, left panel). Expression levels in cells from individual experiments varied between levels observed in A549 and A431 cells (Figure 1E, right panel, Figure S1A-B), which are thought to reflect physiological and supra-physiological levels, respectively (44). Stably-transfected cells were also created with each EGFR TM variant under control of an inducible promoter (45). These cells frequently lost expression or resulted in expression level profiles comparable to transiently transfected cells, however, and experiments with these cells were not pursued.

To test the response of EGFR TM variants to ligand, Western blots probing for phosphorylation at five intracellular EGFR tyrosine residues were performed on lysates from wild-type and variant-expressing cells that were either untreated or treated with EGF (Figure 2A-C). We reasoned that if EGFR TM dimer conformations reflected inactive, active, or activity-modulated states then EGFR variants with TM region substitutions designed to favor or disfavor specific TM dimers would show altered activity. Wild-type EGFR and all variants except Neu (rat ErbB2 V664E) showed reproducible 2.5-5 fold increases in normalized phosphotyrosine band intensities following treatment with EGF. The Neu (rat ErbB2 V664E) TM variant was constitutively phosphorylated at all sites in the absence of EGF, consistent with the ability of this TM variant to drive phosphorylation and tumor formation (46, 47). Addition of a kinase-inactivating substitution (D855N) to both native EGFR and the poly-leucine EGFR TM variant resulted loss of autophosphorylation in response to ligand indicating that EGFR kinase activity is required for receptor phosphorylation in the presence of ligand (Figure S2). Although the phosphorylation levels of all variants except Neu (rat ErbB2 V664E) consistently increased in response to ligand, the normalized levels of increase varied in independent experiments as shown in bar graphs with relatively large standard deviations (Figure S3A-E). In particular, TM regions containing poly-leucine regions often appeared to have smaller increases in phosphorylation in response to ligand, but these differences were within standard deviations of other TM variants. The reproducibility of these observations provides confidence in qualitative interpretations of the TM variant activity, and we conclude that the presence or absence of a GxxxG motif at any position in the EGFR TM is neither required for ligand-dependent EGFR activity nor locks EGFR into an ‘on’ or ‘off’ state. We also note that both strong (Glycophorin A) and weak (Glycophorin A mutant) TM dimers are consistent with ligand-dependent EGFR activity as is a polyleucine TM. The quantitative variability of these results provides insufficient basis for interpreting quantitative differences in TM variant behavior, however.

**Figure 2.**
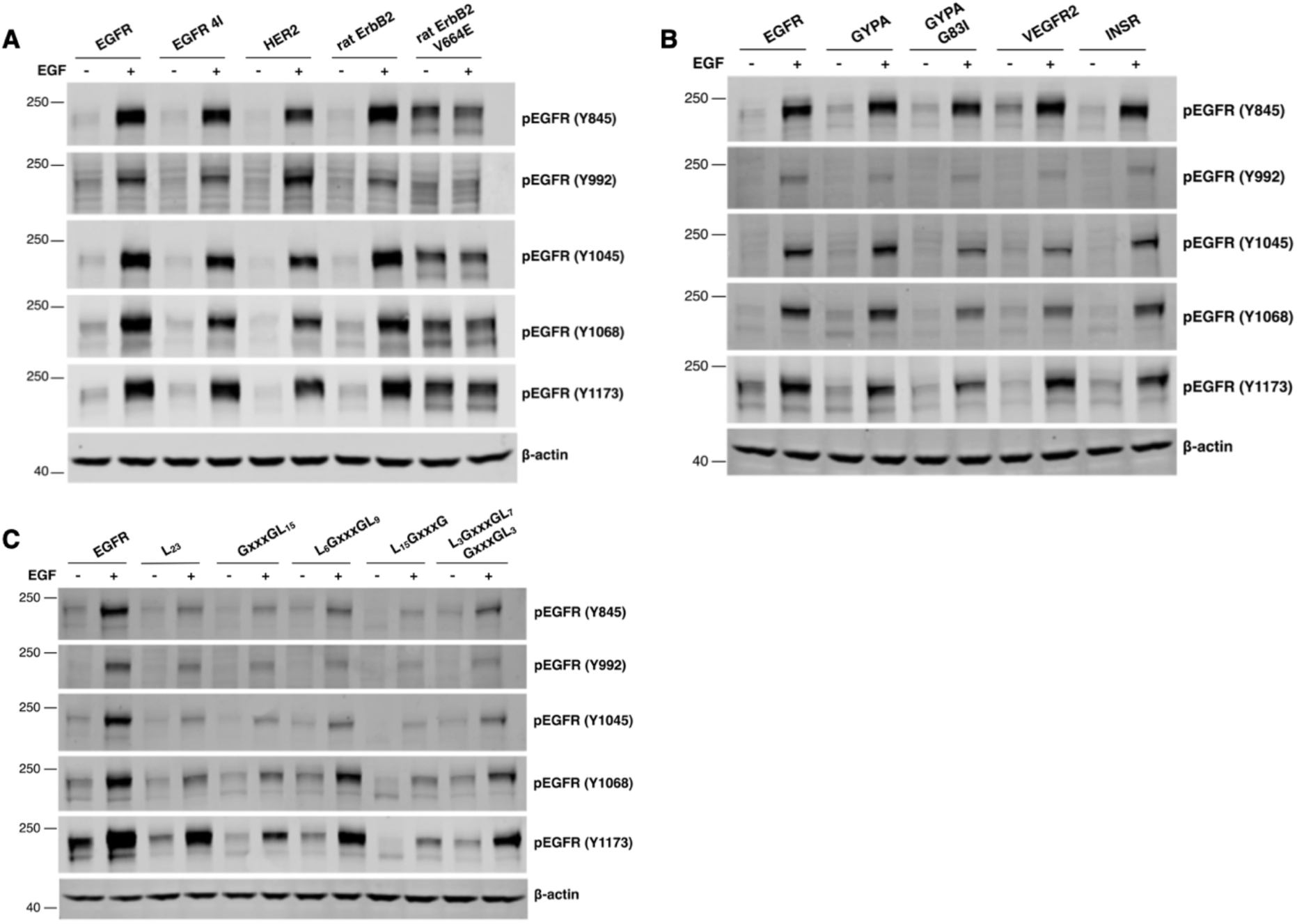
Key phosphorylation sites of all EGFR TM variants are phosphorylated following addition of EGF. (A-C) Western blot analysis of the phosphorylation of EGFR TM variants transiently transfected into CHO cells with and without addition of EGF (100 nM). Phosphorylation was detected using anti-phospho-tyrosine antibodies targeting specific sites in EGFR (pY845, pY992, pY1045, pY1068 and pY1173). β-actin blots are shown as loading controls. Western blots were performed at least three times.

As coupling between receptor and effector phosphorylation is not always present (48), we examined the ability of EGFR TM variants to activate downstream pathway effectors. Western blots probing for phospho-Erk (pErk) and phospho-Akt (pAkt) were performed on lysates from transfected cells treated or untreated with EGF (Figure 3A-C). Stimulation of transfected cells with EGF led to increases in both pErk and pAkt for all TM variants including Neu (rat ErbB2 V664E). Although these increases were reproducible, the magnitude of the responses varied in different experiments leading to high standard deviations, which precluded quantitative interpretation of these results (Figure S4A-B). As it has been suggested that the high-affinity EGFR ligands EGF and TGFα are capable of eliciting distinct cellular responses and that TM dimer conformations may underlie this biased agonism (38), we examined the responses of EGFR TM variants to stimulation with TGFα. Reproducible increases in pErk and pAkt levels comparable to those observed following EGF stimulation were observed in cells transfected with all EGFR TM variants following treatment with TGFα (Figure 4A-C, S5A-C).

**Figure 3.**
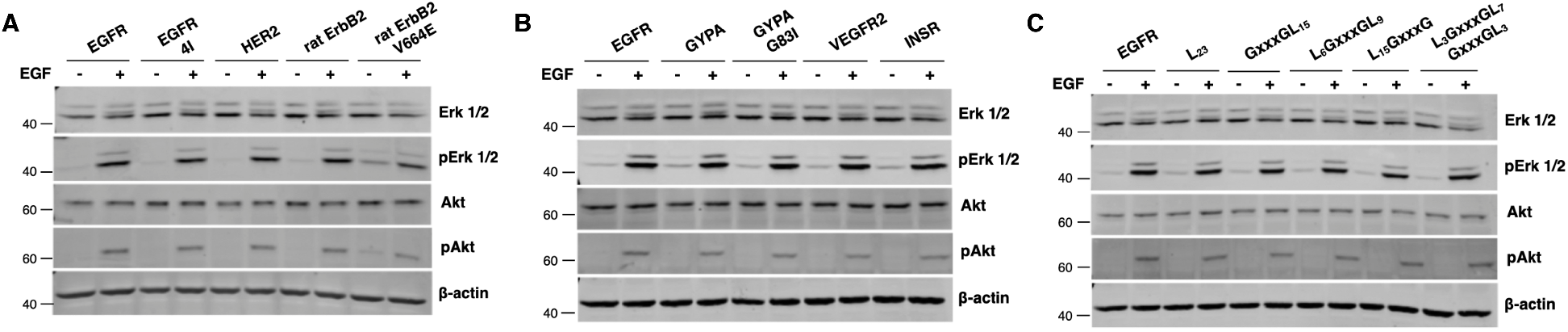
Addition of EGF to all EGFR TM variants leads to phosphorylation of Erk and Akt. (A-C) Western blot analysis of EGF-dependent phosphorylation of Erk and Akt. Expression of Erk and Akt was detected using anti-Erk and anti-Akt (pan) antibodies, and phosphorylation of Erk and Akt was detected with anti-phospho-Erk 1/2 (pErk1/2) and anti-phospho-Akt (pAkt) antibodies, respectively. Experiments were performed in triplicate, and β-actin blots are shown as loading controls.

**Figure 4.**
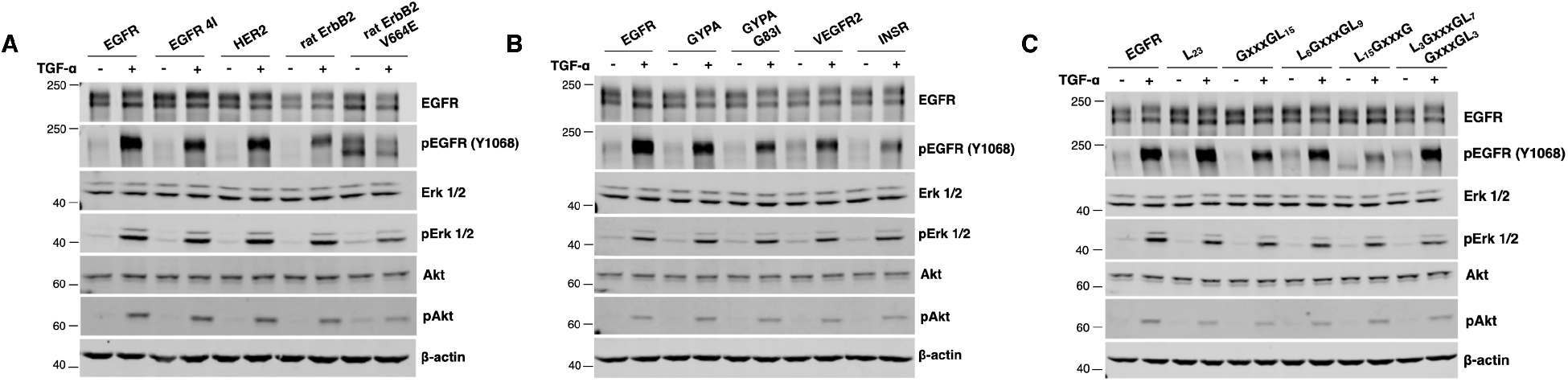
Stimulation of cells with TGFα leads to phosphorylation of Erk, and Akt. (A-C) Western blot analysis of TGFα-dependent phosphorylation of Erk and Akt. Expression of Erk and Akt was detected using anti-Erk and anti-Akt (pan) antibodies, and phosphorylation was detected with anti-phospho-Erk 1/2 (pErk1/2) and anti-phospho-Akt (pAkt) antibodies. Experiments were performed in triplicate, and β-actin blots are shown as loading controls.

## Discussion

Crystal structures of EGFR extracellular and kinase regions have revealed inactive monomeric and active dimeric conformations of these fragments (13–15, 20–24). Specific interactions between EGFR transmembrane regions in active, inactive, and activity-modulated states have been proposed (33, 37, 38), and a dimer interaction mediated by an N-terminal GxxxG motif was observed by NMR for the EGFR TM region in lipid bicelles (34). In contrast, other authors have observed that insertions, deletions, or mutations in the EGFR TM have no detectable effect on EGFR activity (27, 29). We show here by QI-FRET that the EGFR TM region has an intrinsic propensity to self-associate in mammalian cell membranes, consistent with its behavior in TOXCAT assays (25). Failure to observe EGFR TM interactions in detergents likely indicates that confinement to a two-dimensional membrane contributes significantly to observed levels of TM interaction (26, 49). We further show that the EGFR extracellular region reduces association of the TM region in the absence of ligand, but that addition of ligand restores TM proximity. A plausible explanation for this behavior is that EGFR TM separation and association are associated with inactive and active EGFR states, respectively, as has been observed for the Insulin Receptor family (43). If correct, this model implies that the EGFR extracellular region keeps EGFR inactive in the absence of ligand by holding the TM regions apart rather than in an inactive dimer. Further studies are needed, however, to determine if EGFR TM regions indeed remain separated in dimers of full-length EGFR, which are known to form in the absence of ligand (6, 7).

The EGFR TM region contains N-terminal and C-terminal GxxxG motifs that have been proposed to mediate EGFR TM interactions in active and inactive states, respectively, and different EGFR TM dimers found by molecular dynamics simulations have been proposed to influence juxtamembrane region conformations and modulate EGFR signaling outputs (38). We thus generated a panel of thirteen EGFR TM variants that included variants created by eliminating the N-terminal GxxxG motif (4I); substituting the TM region with polyleucine (L_23_); adding back an N-terminal, middle region, or C-terminal GxxxG motif to the polyleucine variant (GxxxGL_15_, L_5_GxxxGL_9_, and L_15_GxxxG, respectively); substituting the TM with strongly and weakly dimerizing Glycophorin A TM variants (GYPA and GYPA G83I, respectively); substituting the TM with the human and rat HER2/ErbB2 and rat Neu TMs (HER2, rat ErbB2, and rat ErbB2 V664E, respectively), and substituting the TM with the Insulin Receptor (INSR) and Vascular Endothelial Growth Factor Receptor (VEGFR2) TMs, which both lack conserved GxxxG motifs.

In Western blot analyses, all variants but the Neu (rat ErbB2 V664E) were reproducibly comparable to native EGFR in their normalized levels of phosphorylation in the absence of ligand and their ability to autophosphorylate and activate Erk and Akt in response to ligand. The Neu (rat ErbB2 V664E) variant was constitutively phosphorylated in the absence of ligand as expected from many previous studies (47, 50). Although qualitatively reproducible, considerable quantitative variability was observed in different Western blot experiments, presumably owing to variations in transfection efficiencies and differing lots of antibodies in replicate experiments among other variables. This variability indicates that our experiments would not reliably detect subtle differences in activity between variants, which may also be affected by high levels of receptor expression in transiently transfected cells.

We nonetheless can conclude that each variant except the Neu variant is capable of remaining inactive in the absence of ligand and activating both itself and downstream effectors in response to ligand. Of particular note, the ability of a polyleucine TM region and TMs with disrupted or specifically placed GxxxG regions to support ligand-dependent receptor and pathway activation implies that multiple types of TM interactions are compatible with ligand-dependent EGFR activity. Our results thus provide no evidence that specific types of TM dimers are required to form inactive or active EGFR dimers and show that many types of EGFR TM association are compatible with receptor activation. We note, however, that our experiments were carried out at relatively high receptor and ligand concentrations and that specific TM motifs may play a role in shaping EGFR activity in low receptor and/or ligand concentrations as has been suggested for cytokine receptors (26, 30).

## Methods

### Plasmid generation

For FRET experiments the extracellular and transmembrane regions of EGFR were cloned into pCDNA3.1(+). A flexible 8-amino acid linker (GSGGSGGS) and fluorescent proteins (eYFP or mCherry) were added to the C-terminal end. For the plasmid containing just the EGFR TM domain, the EGFR signal peptide was maintained, and the extracellular domain was omitted.

For Western blot and immunofluorescence analysis of the variants, DNA encoding each EGFR TM variant was introduced into a previously generated pSSX-EGFR-eYFP expression vector (43). A linear fragment of the initial vector and EGFR coding regions without the transmembrane region was amplified by Polymerase Chain Reaction (PCR) using forward and reverse primers flanking the TM encoding region. DNA segments encoding variant transmembrane regions were then generated in a two-step reaction. Oligonucleotides with 20 bp overlapping regions specific for each transmembrane domain variant were mixed and annealed in a thermal cycler at 94°C for 5 mins then cooled 0.1°C/sec to 5°C below the melting temperature of the primers, kept at the primer melting temperature for 5 mins, and cooled further at 0.1°C/sec to 37°C. The annealed products were incubated for 1h at 37°C with the addition of dNTPs and Klenow fragment, yielding 75bp products encoding each variant TM region. These variant TM-encoding products were then cloned into the pSSX backbone using Gibson assembly. Generation of the kinase inactive variants was achieved using QuikChange PCR (Agilent).

### FRET analysis of membrane-derived vesicles

Cell culture, vesiculation, data collection and analysis were performed as described in (7). Briefly, CHO cells were cultured in 6-well plates in DMEM/F12 + 10% FBS. The medium was changed just prior to transfection. Cells were transfected using a 1:3 (w/w) mixture of plasmid DNA and Lipofectamine 2000, according to the manufacturer’s protocol. Transfected cells were grown for 18-24 hours, then washed twice with 30% PBS and treated with vesiculation buffer (200 mM NaCl, 5 mM KCl, 0.5 mM MgCl_2_, 0.75 mM CaCl_2_, 100 mM bicine pH 8.5) (51). The vesiculation medium was transferred to a chambered coverglass (Lab-Tek II, Nunc, ThermoFisher Scientific), then mounted onto a Nikon Eclipse confocal laser scanning microscope equipped with a 60X water immersion objective lens. Each vesicle was scanned three times: a “donor scan” (488 nm excitation, 500-530 nm filter), an “acceptor scan” (543 nm excitation, 650 nm longpass filter) and a “FRET scan” (488 nm excitation, 565-615 nm filter). The microscope was calibrated using purified fluorescent protein standards of known concentration, which allowed calculation of the 2D concentration of receptor molecules per unit area of the membrane (molecules/□m^2^). The data were fit to a monomer/dimer equilibrium model using GraphPad Prism:

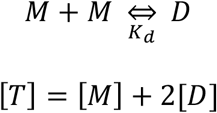

where [M] is the concentration of monomers, [D] is the concentration of dimers, [T] is the total receptor concentration and K_d_. is the dimer dissociation constant. The dimeric fraction (f_D_) within each vesicle is:

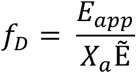

where E_app_ is the apparent FRET efficiency at a given concentration, X_a_ is the fraction of acceptor molecules in a vesicle and 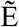 is the FRET value within a dimer (in this case 0.51, the value corresponding to EGFR ECDTM + 100 nM EGF).

### Flow cytometry

CHO-K1 (CCL-61, ATCC), A431 (CRL-1555, ATCC) and A549 (CCL-185, ATCC) cell lines were maintained in complete growth medium (DMEM/F12 supplemented with 10% FBS) and MCF7 (HTB-22, ATCC) cells were maintained in complete growth medium supplemented with 0.01mg/mL insulin. Cells were seeded in 60 mm plates in triplicate, and CHO cells were transfected with 3 μg of expression vector DNA using PEI. 48 hours after seeding, media were removed and cells were detached by incubation with 10 mM EDTA for 15 minutes at 37°C. Cells were spun down and washed three times with cold Flow Wash Buffer (FWB) (PBS, 2% FBS, 2mM EDTA) and incubated with primary antibody (EGFR-D1D4J, Cell Signaling Technology) diluted in FWB for 60 minutes on ice. Cells where then washed three times with FWB and incubated with secondary antibody (Alexa Fluor 555-A21428, Invitrogen) diluted in FWB for 60 minutes in the dark, on ice. Cells were finally washed three times and resuspended in FWB, passed through a cell strainer (Stemcell Technologies) and analyzed using flow cytometry. Fluorescence analysis was performed with a FACScan cytometer (LSRFortessa SORP Flow cytometer) (BD Biosciences) using a 561 nm excitation laser. Sample acquisition and analysis were done using FACSDiva v9 and FlowJo software, respectively.

### Cell culture and cell lysis

CHO cells were maintained at 37°C in DMEM/F12 medium supplemented with 10% FBS. Cells were seeded in 6-well plates and transfected with 1.5 μg of each variant-encoding plasmid using PEI. After 24 hours, the cells were washed three times with warm PBS and serum starved for 3 hours in Ham’s F-12 medium supplemented with 1 mg/mL BSA. Starvation was followed by stimulation with EGF or TGF-α at 100 nM or 25 nM final concentration, respectively, for 5 minutes at 37°C. Cells were then washed three times with cold PBS and lysed in cold RIPA buffer (50 mM Tris pH 8.0, 150 mM NaCl, 1% v/v NP40, 0.5% w/v sodium deoxycholate, 0.1% w/v SDS) supplemented with 1 mM activated Na_3_VO_4_, one protease inhibitor mini-tablet (Thermo Scientific) and Benzonase nuclease (Millipore). Cell lysis was performed for 30 minutes at 4°C under gentle rocking.

### Western blotting

Cell lysates were clarified by centrifugation, and total protein concentration was determined using a BCA assay (Pierce BCA protein assay kit, Thermo Scientific). Lysates were normalized for total protein concentration, mixed and boiled with 5X sample loading buffer (Biolegend) and loaded on 4-12% Tris-glycine gels (Invitrogen). Protein bands were transferred to nitrocellulose membranes using the iBlot® Dry Blotting System. Membranes were blocked with 5% non-fat milk in TBS for 1 hour at room temperature and subsequently incubated with primary antibodies for the detection of EGFR (D38B1), EGFR-pY1068, EGFR-pY845, EGFR-pY992, EGFR-pY1045, EGFR-pY1173 (53A5), Erk 1/2 (137F5), pErk 1/2, Akt (pan) (11E7) and pAkt (D9E) for 16 hours at 4°C. Membranes were then washed three times with TBS (Tris-buffered saline; 50 mM Tris pH 7.5, 150 mM NaCl), incubated with the appropriate IRDye secondary antibody (Goat anti-rabbit 680RD, Licor) and imaged using the Odyssey CLx Imaging System (LICOR). Western blot images were analyzed and quantified with Image Studio Lite software. All primary antibodies were purchased from Cell Signaling Technology.

### Immunofluorescence

CHO cells were seeded in glass coverslips, grown in DMEM/F12 medium supplemented with 10% FBS and transfected with 1.5 μg DNA in the pSSX vector using PEI. 24 hours post transfection the coverslips were washed with PBS, fixed in 4% paraformaldehyde (PFA) for 20 minutes at room temperature and blocked with PBTG buffer (PBS, 1% BSA, 1M glycine, 0.1% Triton X-100) or PBG buffer (no Triton X-100) when permeabilization was not required. Samples were incubated with primary antibodies (EGFR-D1D4J, Cell Signaling Technology and/or pEGFR-MA5-15199, Invitrogen) diluted in 1% BSA in PBS for 1 hour at room temperature in a dark humid chamber. Cells where then washed with PBS and incubated with secondary antibodies (Alexa Fluor 555-A31572, Alexa Fluor 555-A21428, Alexa Fluor 633-A21052 or Alexa Fluor 488 coupled to wheat germ agglutinin, Invitrogen) for 30 minutes. After secondary antibody removal, the coverslips were incubated with 1 μg/mL DAPI stain for 5 minutes, washed and mounted on glass slides using Prolong Gold anti-fade reagent (Invitrogen). Images were taken with a Zeiss 710 Laser Scanning Confocal (ZeissCF) microscope and analyzed using Fiji software.

A431, A549 and MCF7 cells were maintained as described previously and stained according to the above protocol. Images were acquired in the Zeiss 710 Laser Scanning Confocal (ZeissCF) microscope using a plan-apochromat 63x/1.4 Oil DIC M27 objective, at same conditions of laser power, gain and pixel dwell time. Images of 8 bpp size of 1024×1024 were analyzed using the open-source software package Image J to get the plot profile across cells with different levels of expression within a single image.

This article contains supporting information.

## Acknowledgments

We thank Kevin Dalby and Sean Burke for helpful discussions.

## Funding and additional information

This work was supported by National Institutes of Health Grant 5R01GM099321 and Cancer Prevention Research Institute of Texas (CPRIT) Grant RR160023 (DJL) and National Institutes of Health Grant R01 GM068619 (KH). The content is solely the responsibility of the authors and does not necessarily represent the official views of the National Institutes of Health.

## Ethics Statement

No animals were used in the reported work.

## Data Availability Statement

No new crystallographic or sequence data are reported in this manuscript. The authors agree to make any materials, data, and associated protocols available upon request.

## Abbreviations

EGFR: epidermal growth factor receptor;
RTK: receptor tyrosine kinase;
HER: human epidermal growth factor receptor;
TM: transmembrane domain;
CHO: Chinese hamster ovary;
fp: fluorescent protein;
QI-FRET: quantitative imaging FRET.

