## Supplemental Figures for "Activity of EGFR transmembrane region variants indicates specific transmembrane dimers are not required for EGFR activity"

**SUPPORTING INFORMATION**

Foteini Bartzoka[1], Monica Gonzalez-Magaldi[1], Patrick O. Byrne[1], Nicole I. Callery[1], Kalina Hristova[2], and Daniel J. Leahy[1]*

**Table S1.** Kd values as measured by QI-FRET.

| Proteins | [EGF] (nM) | Kd (molecules/μm$^2$) | (95% CI) |
|---|---|---|---|
| EGFR-ECDTM-fp | 0 | 3105 | (2054-4900) |
| EGFR-ECDTM-fp | 100 | <1 | (N.D.) |
| EGFR-TM-fp | 0 | 57 | (29-97) |

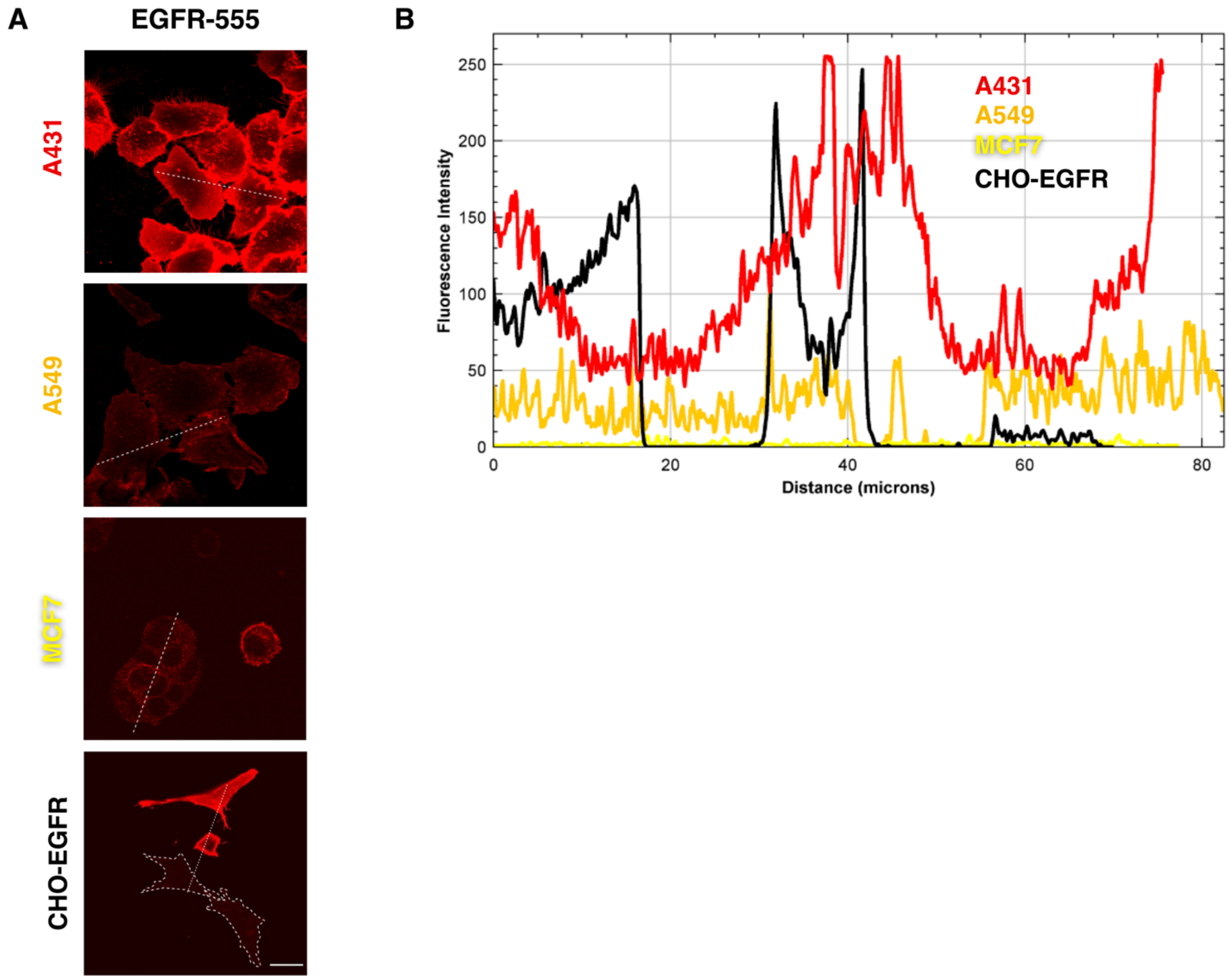


**Figure S1. Levels of EGFR expression quantified by confocal microscopy.** A) Immunofluorescence images of four different cell lines (A431, A594, MCF7 and EGFR-transfected CHO cells) acquired using an antibody targeting the extracellular region of EGFR on fixed non-permeabilized cells. B) Two-dimensional representation of the intensity of pixels along the dotted line (distance in microns) for each image shown on panel A. Scale bar represents 20 μm.

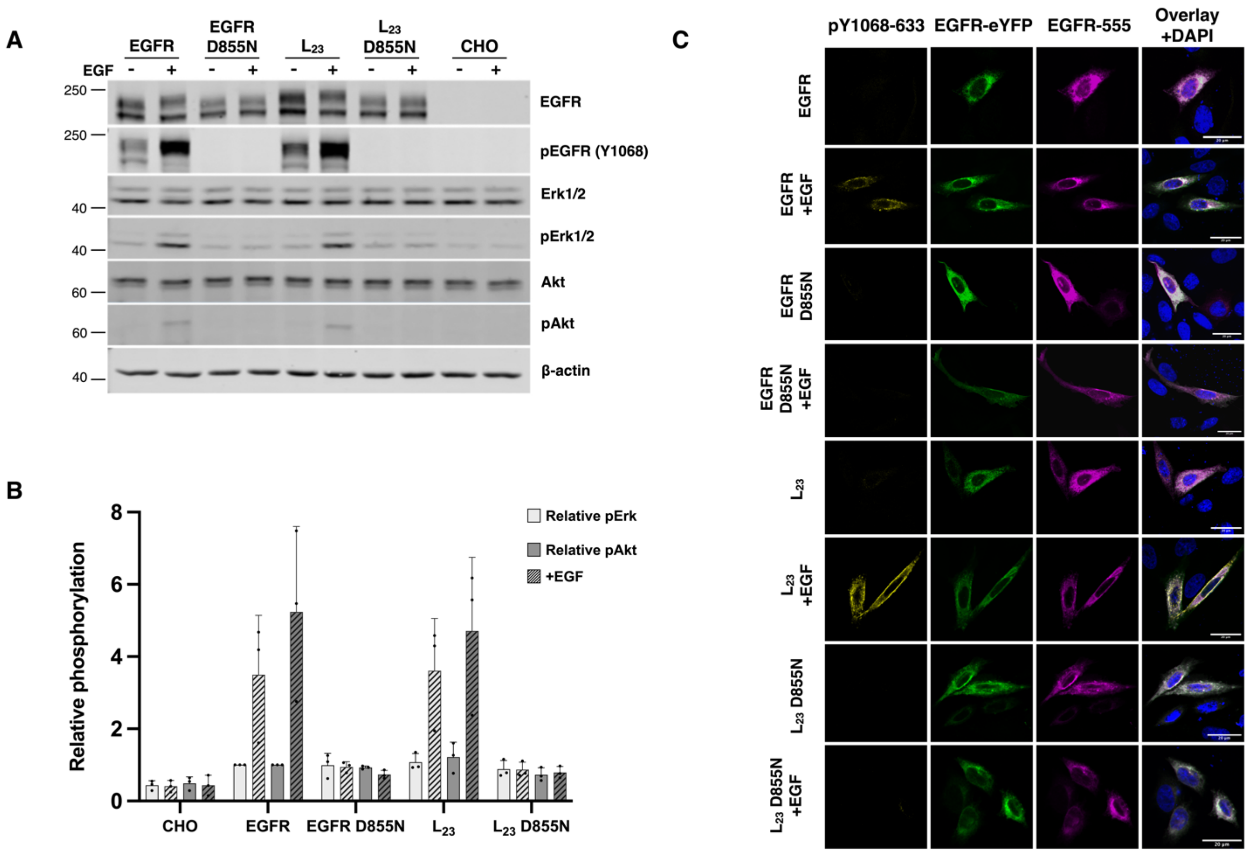


**Figure S2. Kinase activity required for EGFR $L_{23}$ variant and downstream effectors to become phosphorylated in response to EGF.** A) Western blot analysis of the expression and phosphorylation of EGFR, Erk, and Akt in transiently transfected CHO cells without and with EGF (100nM) stimulation. Untransfected CHO cells (CHO) are shown as negative control. Western blot experiments were performed in triplicate, and a β-actin blot is shown as loading control. (B) Quantification of the increase in phosphorylation of Erk and Akt after normalization to Erk and Akt expression levels. These ratios were then normalized to the comparable value for wild type EGFR without EGF. Dots represent three independent experiments, and standard deviations are shown. (C) Confocal microscopy images of transiently transfected permeabilized CHO cells. Receptor expression is shown in purple and green and pY1068 is shown in yellow. Scale bar represents 20 µm. Imaging was performed in triplicate and representative images are shown.

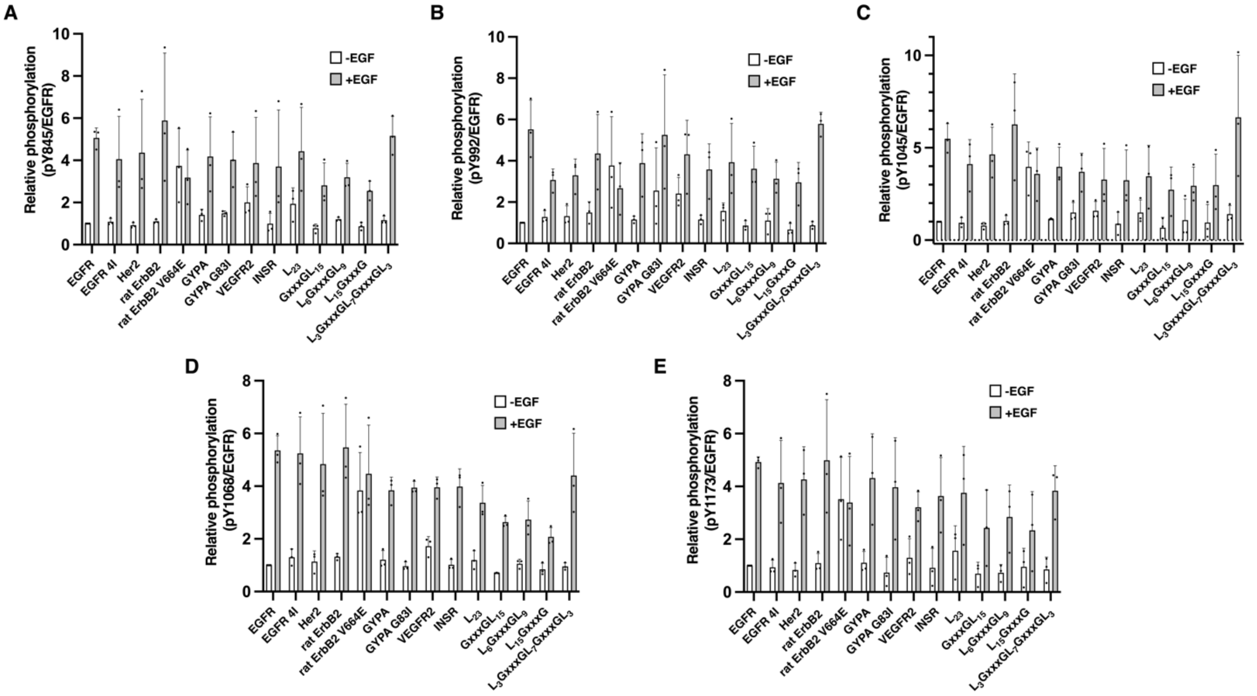


**Figure S3. Quantification of the increase in EGFR phosphorylation upon EGF stimulation.** (A-E) The intensity of tyrosine phosphorylation at each site was normalized to the respective receptor expression, and this ratio further normalized to the comparable ratio for wild type EGFR in the absence of EGF. Cells were stimulated with 100 nM EGF. Dots represent values from three independent experiments and standard deviations are shown.

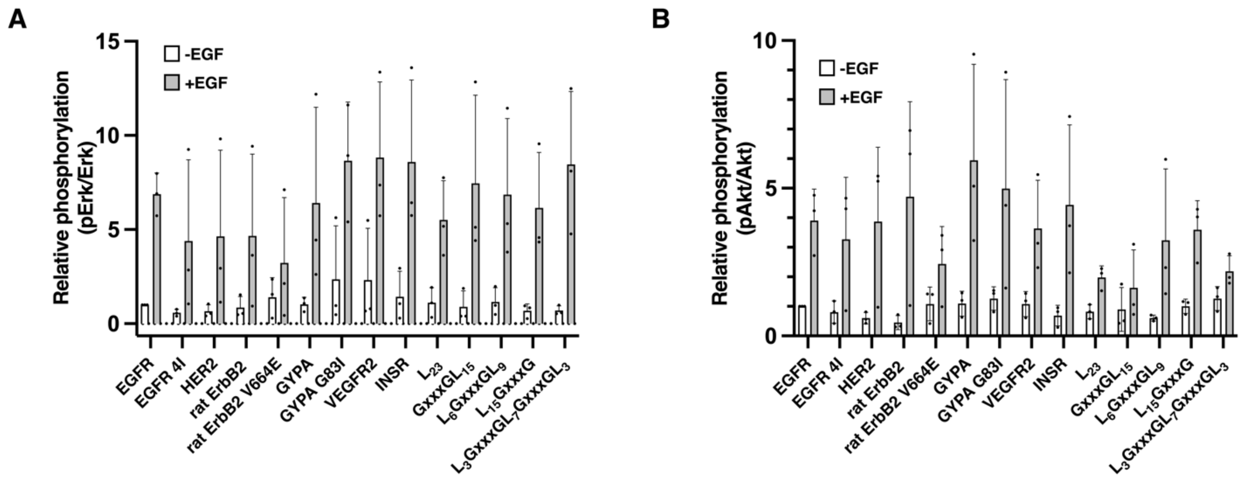


**Figure S4**. **Quantification of the increase in phosphorylation of Erk and Akt.** (A-C) Intensities of pErk and pAkt bands were normalized to Erk and Akt expression levels, respectively, and these values normalized to the comparable ratio for wild type EGFR without EGF. Cells were stimulated with 100 nM EGF. Dots represent three independent experiments and standard deviations are shown.

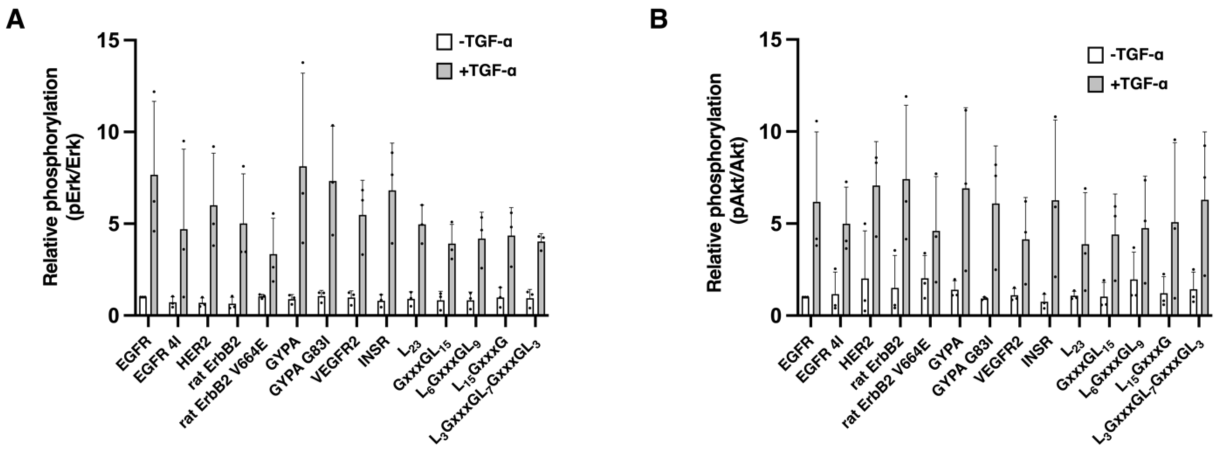


**Figure S5. Quantification of the increase in phosphorylation of Erk and Akt upon TGF-α stimulation.** (A-C) Intensities of pErk and pAkt bands were normalized to Erk and Akt expression levels. Erk and Akt phosphorylation levels were further normalized to that of wild type EGFR without TGF-α. Cells were stimulated with 25 nM TGF-α. Dots represent three independent experiments and standard deviations are shown.
